# A novel PCR method directly quantifies sequence features that block primer extension

**DOI:** 10.1101/325308

**Authors:** Richard M. Cawthon

**Affiliations:** Dept. of Human Genetics, University of Utah

**Keywords:** quantitative PCR, genomics, epigenetics, DNA methylation, DNA damage, aging

## Abstract

Many quantitative polymerase chain reaction (qPCR) assays indirectly quantify a target of interest as a drop in amplification signal relative to a larger total signal. The targeted sequence features include: DNA strand breaks introduced by enzymes that cleave at specific sequences or modifications, or caused by ionizing radiation or other DNA-damaging agents; DNA damage that does not break the DNA but blocks the DNA polymerase; and deletions of various sizes that prevent one or both primers from annealing. All of these loss of signal (a.k.a. negative) assays share the drawback of low sensitivity, as compared to positive assays, which generate an amplification signal that directly reflects the copy numbers of the sequence feature. Here we present a novel qPCR strategy that converts all of the above negative assays into positive ones. A mixture of three primers is added to genomic DNA, one primer pair that targets a sequence for strand-specific PCR, and a third, longer primer that prevents initiation of that PCR, by annealing at high temperature and extending across the target sequence, rendering it doublestranded and inaccessible for priming. Any sequence feature that blocks the third primer’s annealing or extension, while leaving the target sequence intact, allows the PCR to proceed and quantify the copy number of the sequence feature. This Primer Extension Blockade Enabled qPCR (PEBE-qPCR) method will facilitate many high-throughput, low-cost qPCR-based investigations in biology and medicine. As a first example, we present direct qPCR of the unmethylated allele at a MspI/HpaII site in the promoter of the human 45S rDNA gene.

## Background

Archived purified human DNA samples from healthy middle-aged and older adults, accompanied by long-term follow-up survival, cause of death, and cancer incidence data for the donors, are invaluable resources for aging research. In addition to the inherited genotypes that can be scored and tested for association with healthspan and lifespan, many additional sequence features can be detected and quantified in archived DNA. The methylation states at hundreds of loci change with age, at different rates in different people, and “DNA methylation age” based on individual CpG dinucleotide sites predicts life expectancy [1]. Methylation at the promoters of LINE-1 retroelements declines with age, varies between age-matched individuals, and is associated with differences in all-cause mortality [2-3]. Senescent cells accumulate with age in many tissues and can be detected and quantified by their specific methylation signatures [4]. DNA damage of several types accumulates during aging, including oxidative damage [5], double-strand breaks [6], large deletions [7], and point mutations [7-8]. High-throughput, low-cost assays to quantify each of these classes of genomic instability will speed research into the causes of aging-related diseases and disabilities and the development of interventions to extend the human healthspan and lifespan. Here we present a novel qPCR strategy allowing high-throughput, low-cost direct quantification of many different DNA sequence features relevant to biology and medicine.

## Results

### Primer Extension Blockade Enabled Quantitative PCR (PEBE-qPCR)

Figure 1 explains the basic method. A list of sequence features that, in principle, should be quantifiable by this approach is included (Fig. 1C), which is not intended to be exhaustive.

**Figure 1.**
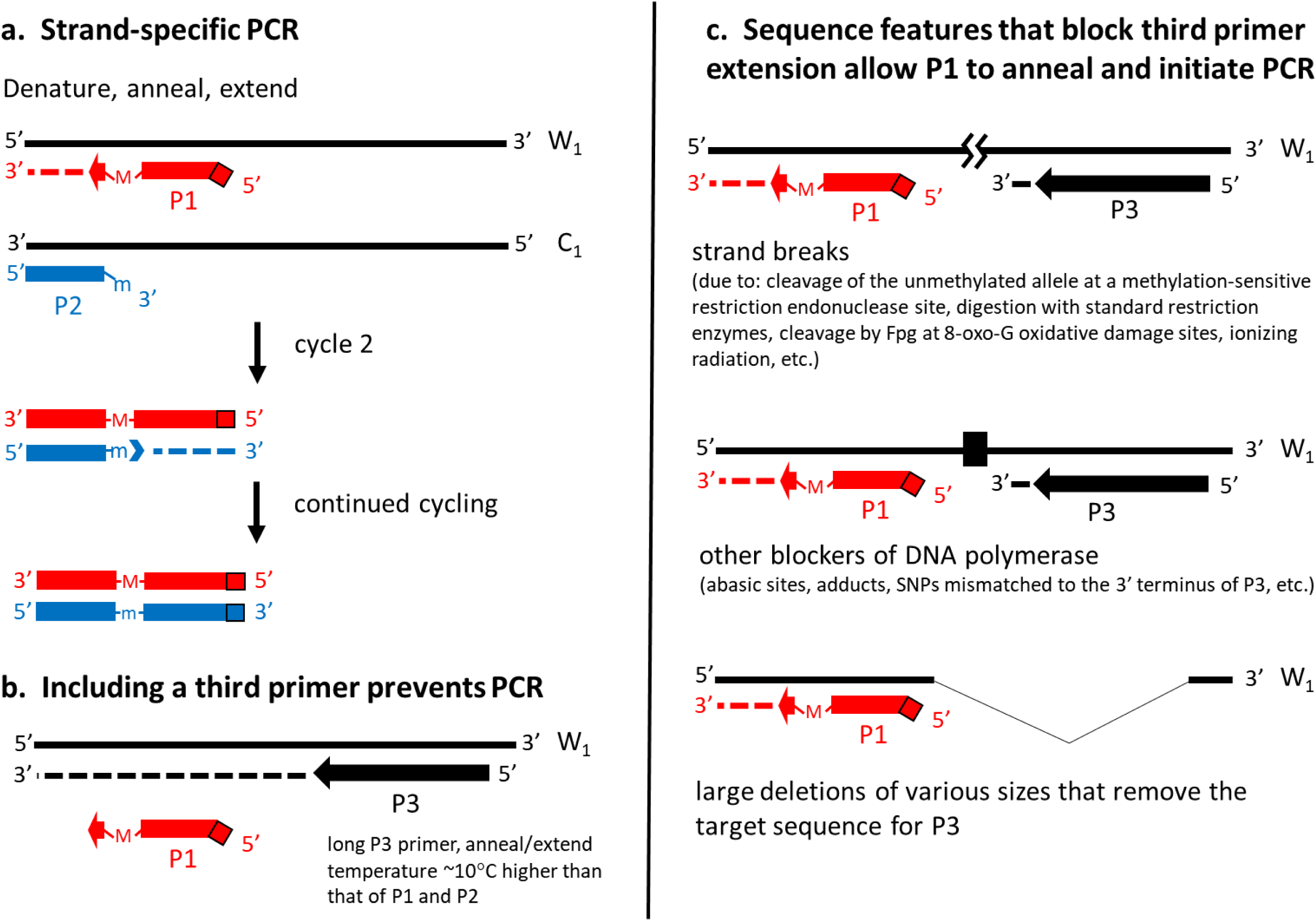
Direct quantification of sequence features that block primer extension. **a.** Strand-specific PCR. Primer P1, intentionally mutated (M) at the third base from its 3’ end and at the last two bases at its 5’ end, anneals and extends along the W_1_ DNA strand. Primer P2 anneals to the C_1_ DNA strand, upstream of and overlapping by 3 bases primer Pi’s target sequence. The mutation (m) at the 3’ terminus of P2 prevents it from extending along C_1_. Because m is complementary to M, in cycle 2 primer P2 is able to prime DNA synthesis along the P1 extension product. Thermal cycling and exponential amplification ensue. **b.** Including a third primer prevents PCR. Annealing and extending P3 at high temperature blocks hybridization of Pi, blocking PCR. (The mutations at the 3’ ends of P2 and its extension product ensure they cannot prime either native DNA or the P3 extension product.) **c.** DNA sequence features that block P3 extension allow Pi to anneal and extend, initiating PCR.

### PEBE-qPCR design for direct quantification of the unmethylated allele at a HpaII / MspI restriction site (5’-CCGG-3’) in the human 45S rDNA promoter

The 45S rDNA occurs as tandem repeats on the p-arms of the five acrocentric chromosomes (Chromosomes 13, 14, 15, 21, and 22). Its copy numbers are dynamic and vary widely between people, with average copy numbers ranging from approximately 60 to > 800 per cell [9]. Lower rRNA expression levels are associated with longer lifespans, both within and across species [10]. The 45S rDNA promoter is mainly methylated at CpG dinucleotides when transcriptionally silent, and mainly unmethylated at those sites when actively transcribed [11]. It is possible that the copy number per cell of 45S rDNA promoters that are unmethylated at a single CpG site will correlate well with rRNA transcription rates, and therefore be a clinically important biomarker.

Figure 2 shows the 45S rDNA promoter sequence segment (bases −132 to −59) and MspI/HpaII site (5’-CCGG-3’, from −87 to −84) that we targeted for analysis, and the sequences of the three primers used, rpdd, rpd, and rpux. Note that for this assay, in addition to introducing a mutation at the 3’ terminal base of primer rpux (corresponding to the “m” mutation of primer P2 in Fig. 1), we introduced a second mutation in rpux, at the fourth nucleotide position from the 3’ end, which further helps in stopping rpux from priming either native DNA or the extension product of primer rpdd. (This strategy for increasing the specificity of primers used in PCR is derived from the allele-specific PCR literature [12-13].)

**Figure 2.**
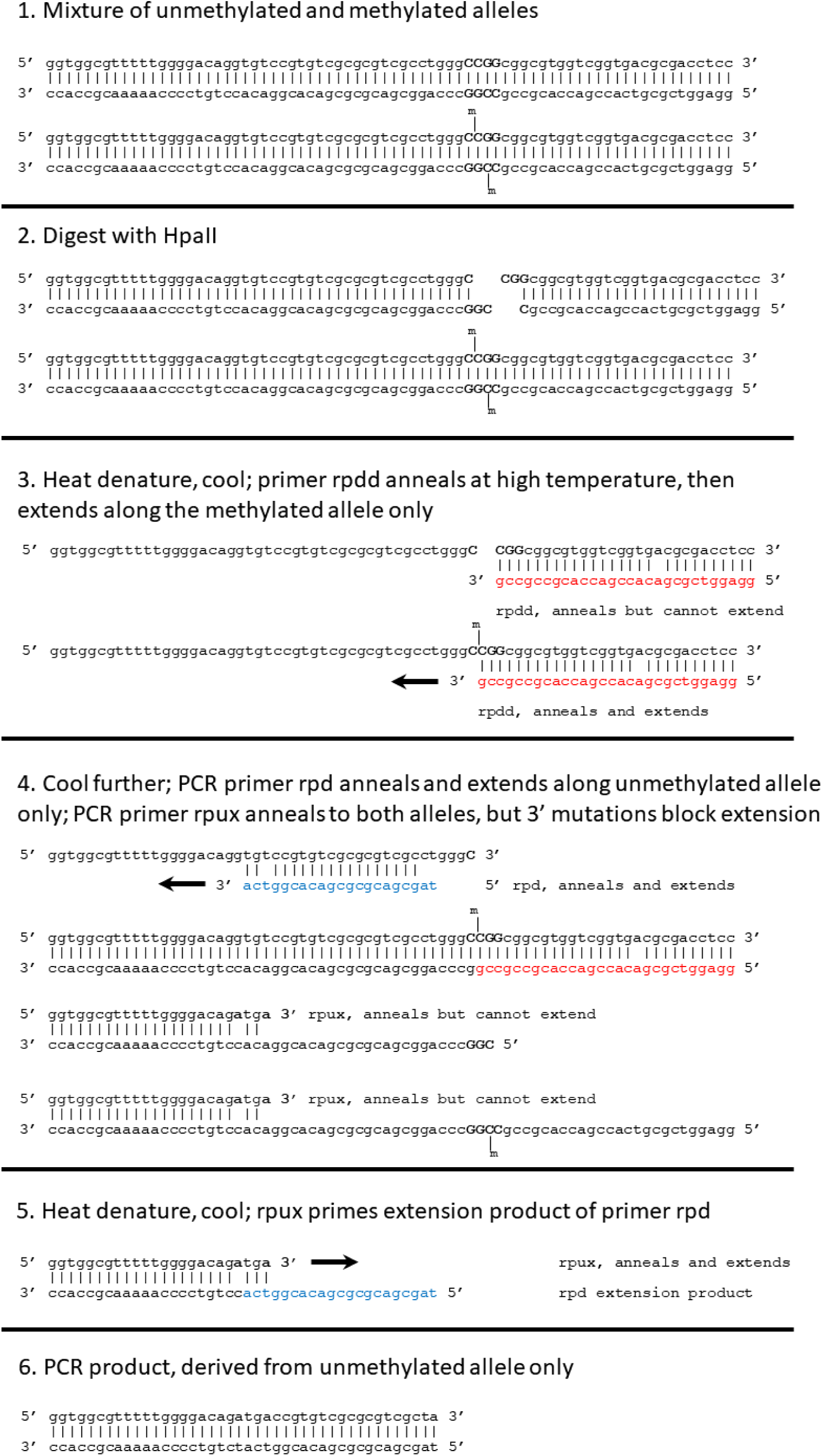
Primer Extension Blockade Enabled qPCR design for the unmethylated allele at the −86 −85 CpG dinucleotide in the human 45S rDNA promoter. All three oligonucleotide primers, rpdd, rpd, and rpux, and the methylation-sensitive restriction endonuclease Hpall, which cuts at unmethylated 5’-CCGG-3’ sites, are combined with PCR master mix, hot-start DNA polymerase, and the DNA sample. The PCR tubes or plates are then sealed, and the restriction digestion and amplification steps carried out on a programmable thermal cycler with real-time fluorescence detection capability. Details are provided in the Methods section.

### Quantitative PCR from the target sequence upstream of the cleaved 5’-CCGG-3’ site, and suppression of PCR when the 5’-CCGG-3’ site is intact and primer rpdd is included

As is typical for many promoters, this segment of the human 45S rDNA promoter has a GC-content greater than 70%, making amplification by PCR difficult. To reduce the formation of secondary structures that might interfere with primer binding, we included 1M betaine to decrease the strength of GC base-pairing [14] and kept all thermal cycling temperatures at 71 °C or higher.

Our goal was to optimize rpux vs. rpd amplification of the primer pair’s target sequence in genomic copies cleaved at the MspI/HpaII restriction site, while maximally suppressing amplification from that same target sequence in genomic copies in which the MspI/HpaII site remained intact.

Figure 3a shows the thermal profile and cycling used. New England BioLabs (NEB) restriction enzymes MspI and HpaII are listed by NEB as Time-Saver™ enzymes, capable of complete digestion in 5-15 minutes at their recommended incubation temperature of 37°C. The slow ramp from 83°C to 81 °C, 30 seconds at 81 °C, and additional slow ramp from 81 °C to 71 °C were designed to allow full annealing and extension of primer rpdd prior to the annealing and extension of primer rpd.

**Figure 3.**
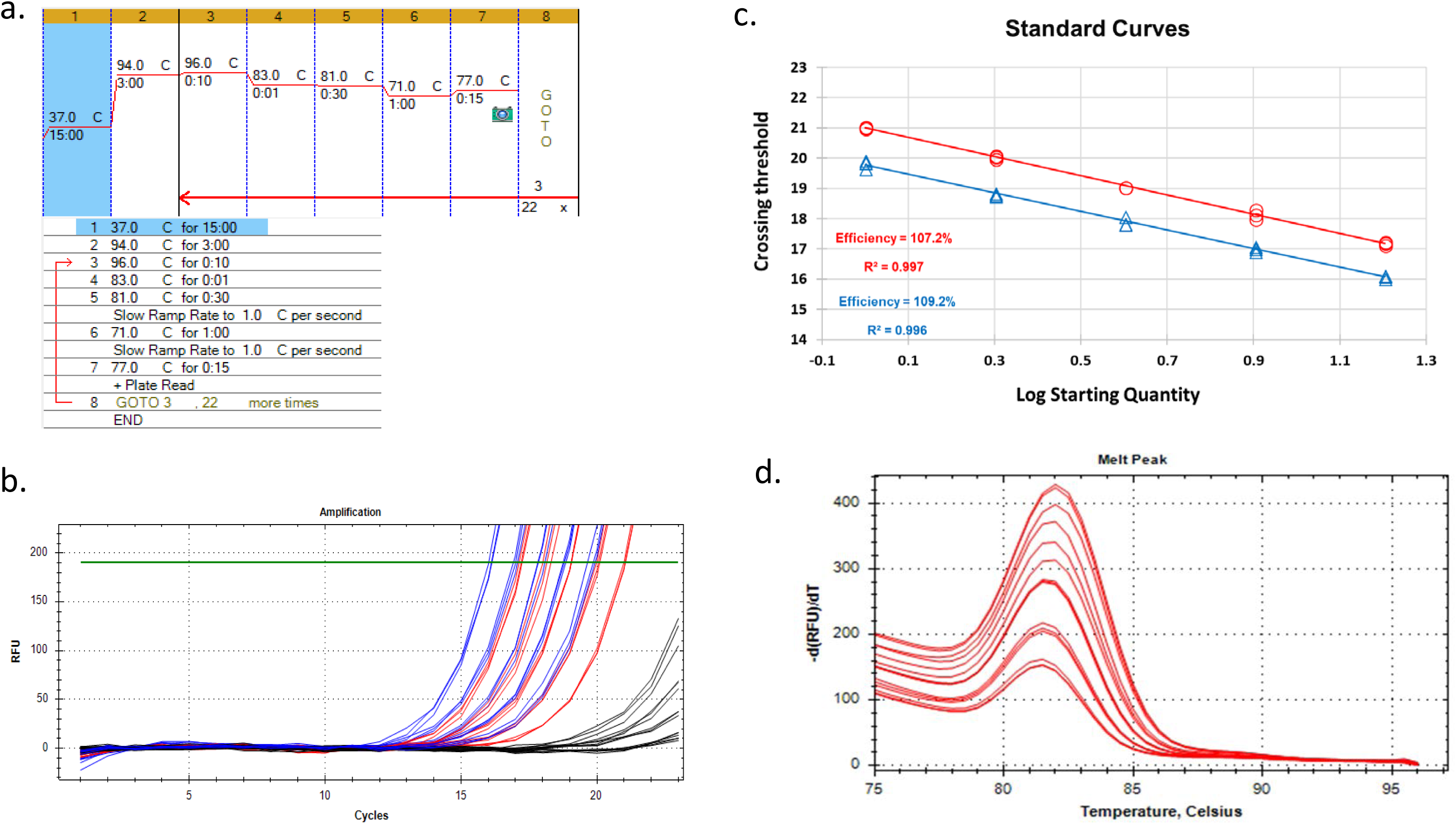
Quantitative PCR of the human 45S rDNA promoter. All reactions received the rpdd, rpd, and rpux primers. Reactions also received either restriction endonuclease Hpall, which cuts unmethylated, but not methylated, 5’-CCGG-3’ sites; restriction endonuclease MspI, which cuts both unmethylated and methylated 5’-CCGG-3’ sites; or no restriction enzyme. Input amounts of the serially diluted DNA per 10 μ.l reaction were 1, 2, 4, 8, and 16 ng of genomic DNA. **a.** Thermal profile and cycling. **b.** Amplification curves for rDNA promoters that are unmethylated at the −86 −85 CpG dinucleotide (red curves, DNA cut with Hpall), for total rDNA promoter copies (blue curves, DNA cut with MspI), and for control samples not treated with restriction enzymes (black curves). **c**. Standard Curves for the same amplification data. Unmethylated allele, red circles and linear regression line; total rDNA promoter copies, blue triangles and linear regression line. **d.** Melt peaks of the PCR products amplified from the unmethylated allele (Hpall treated DNAs). (The melting profiles of the PCR products amplified from the MspI treated DNAs were indistinguishable from these.)

The blue curves in Figure 3b show that robust amplification was achieved in reactions receiving MspI, which cleaves at 5’-CCGG-3’ sites regardless of their methylation state. As expected, including or omitting the third primer, rpdd, had no effect on amplification from MspI treated samples, since the DNA break introduced into all genomic copies by the restriction enzyme blocks rpdd from extending across the target sequence.

Amplification was strongly suppressed (delayed approximately 7 cycles, relative to the blue curves) in reactions omitting restriction enzymes but containing the rpdd primer (black curves in Fig. 3b). This is also as expected, since in undigested samples, rpdd should extend across both the MspI/HpaII site and the target sequence in all genomic copies, preventing rpd from initiating the rpux + rpd PCR.

No amplification was observed in either the No Template Control reactions (those receiving no genomic DNA), or in controls receiving genomic DNA plus the rpux and rpdd primers, but no rpd primer.

### Cutting the 5’-CCGG-3’ site with HpaII followed by primer rpdd extension enables direct and specific qPCR of the unmethylated allele

The restriction endonuclease HpaII produces double-strand breaks at unmethylated 5’-CCGG-3’ sites, leaving methylated 5’-CCGG-3’ sites intact. Therefore, PCR with rpux + rpd in the presence of rpdd, of genomic DNA that has been cut with HpaII, should amplify from the cut genomic copies (i.e. all those that are unmethylated at the 5’-CCGG-3’ site), while suppressing amplification from the uncut genomic copies (i.e. those that are methylated at the 5’-CCGG-3’ site). The amplification curves for these reactions are shown in red in Fig. 3b. These curves’ crossing threshold values (Cts) were collected at cycle numbers when otherwise identical but undigested reactions’ amplification signals were at baseline (black curves in Fig. 3b), indicating that amplification from the uncut methylated genomic copies remaining after HpaII digestion is adequately suppressed by rpdd extension.

Interestingly, the reactions in which amplification was suppressed by rpdd extension in the absence of restriction digestion definitely showed non-zero amplification, proportional to the amount of input DNA, and delayed by approximately six cycles, relative to the amplification of the HpaII treated samples. We were unable to eliminate this low level of amplification by various tweaks to the reaction conditions. This amplification could be due to a small fraction of the genomic copies in this reference pooled DNA sample (derived from eight healthy individuals aged 65 years or older) having breaks in the DNA between the binding sites of the rpdd and rpd primers, and/or deletions or other types of DNA damage preventing rpdd annealing and extension, but still allowing rpux + rpd PCR further upstream. DNA replication *in vivo* through ribosomal DNA tandem repeats is known to be prone to replication fork stalling [15]; unrepaired DNA strand breaks arising during these processes and remaining in the extracted DNA may be contributing to the low level amplification signals we observe in the reactions that receive no restriction enzymes.

At the end of the PCR, a single melt peak (T_m_ approximately 82°C) was detected (Fig. 3d), which was indistinguishable between MspI and HpaII treated samples.

Since amplification after cutting with HpaII (red curves in Fig. 3b) quantifies the unmethylated copies of the 45S rDNA promoter, and amplification after cutting with MspI (blue curves) quantifies the promoter’s total copy number, the difference in crossing thresholds (ΔC_t_) for blue vs. red, at each DNA input amount, allows us to directly measure the fraction of rDNA promoter copies that is unmethylated at this CpG site, and to calculate the fraction that is methylated.

Fig. 3c shows standard curves for amplification from the 2-fold serial dilution of the DNA sample cut with either MspI (in blue) or HpaII (in red). The Cts for the MspI treated samples occurred approximately 1.17 cycles earlier than those collected for the HpaII treated samples. Therefore, the total number of rDNA promoter copies in the sample is approximately 2^ΔCt^ = 2.25 times the number of rDNA promoter copies that are unmethylated at the CpG site. Therefore, in this pooled DNA sample from eight individuals, the unmethylated fraction is approximately 0.44 and the methylated fraction is approximately 0.56.

Quantitative PCR assays for a single copy gene (such as the albumin gene, ALB, or the betaglobin gene, HBB) provide an amplification signal that serves as a proxy for cell count. Therefore, by performing rDNA promoter qPCR and single copy gene qPCR on the same DNA samples, and normalizing the first signal to the second signal, one can perform relative quantification of the copy number of 45S rDNA promoters that are unmethylated at the CpG site *per cell*, in DNAs digested with HpaII; and relative quantification of the total copy number of 45S rDNA promoters per cell, when the DNA is digested with MspI. We hypothesize that the number of unmethylated copies per cell will have clinical importance, since that measure is expected to directly reflect the rate of synthesis of ribosomal RNA in the cells from which the DNA was extracted. (Since the total copy number of the 45S rDNA tandem repeats in humans vary over more than a 10-fold range, from approximately 60 copies per cell to more than 800 copies per cell [9], people with identical rDNA promoter unmethylated fractions may nevertheless differ greatly in the number of unmethylated copies of the rDNA promoter per cell.)

We are now using this qPCR assay to determine whether individual differences in the copy number of the unmethylated 45S rDNA promoter per cell are associated with differences in overall survival, cause-specific mortality, and cancer incidence in 147 subjects from the Utah CEPH [16] grandparent generation (manuscript in preparation).

## Discussion

This novel PCR strategy, Primer Extension Blockade Enabled Quantitative PCR, allows direct quantification of many different DNA sequence features relevant to biological and medical research. In principle, the copy numbers of all of the following can be directly measured by this method: 1) the unmethylated allele at any methylation-sensitive restriction site; 2) somatic mutations that create restriction endonuclease recognition sites (e.g. the A3243G mutation in mtDNA); 3) DNA segments containing one or more 8-oxo-guanine sites due to oxidative damage, cleavable by Fpg (formamidopyrimidine [fapy]-DNA glycosylase); 4) single-stranded or double-stranded DNA breaks due to ionizing radiation and other causes; 5) DNA segments with one or more sites that are abasic or are damaged in other ways that block DNA synthesis by DNA polymerase; and 6) large deletions of various sizes (such as those that accumulate in mtDNA in skeletal muscle, heart, and brain during normal aging). This is not meant to be a complete list; additional applications of the method are also possible.

In the rDNA promoter assay presented above, the rpdd primer extension that prevents rpd primer annealing (corresponding to the P3 primer extension preventing P1 primer annealing in Fig. 1) occurs in each cycle; however, in principle, P3 primer extension can be limited to a single incubation step and still have its full effect of preventing P1 + P2 amplification from the genomic copies that lack the P3 extension-blocking sequence feature(s) of interest. For example, one can use a P1 primer with a templated sequence (the portion that hybridizes to the native genomic template) that anneals at ∼ 60°C, plus a 20-30 base long GC-rich 5’ tagging sequence, in combination with a long P2 primer with a high annealing temperature. This alternative design preserves the first three steps of the original protocol: full denaturation of the DNA (∼95°C); followed by annealing and extending primer P3 at a temperature (∼80°C) too high for the templated portion of primer P1 to anneal; followed by annealing and extending primer P1 at ∼60°C along the remaining available genomic copies. However, after these three incubations, thermal cycling simply alternates between full denaturation (∼95°C) and a high (∼80°C) anneal/extend/signal acquisition temperature for the rest of the run. In this modified protocol, P1 only primes DNA synthesis along the native genomic DNA once (instead of once each cycle, as in the original protocol).

By using PEBE-qPCR in combination with long PCR [17], it may be possible to obtain a measure of the total load of DNA damage of various types in a single high-throughput, low-cost assay. For example, a P3 primer could be annealed to the mtDNA region that is prone to large deletions, and then extended, under conditions optimized for long PCR, along a several kilobase stretch of the mitochondrial DNA. Using the modified protocol described in the previous paragraph, this long duration (e.g. 12 minute) extension need be done only once, followed by rapid thermal cycling with primers P1 and P2. In principle, with this design P1 + P2 should not amplify from intact, damage-free mtDNA copies, but should amplify from mtDNA copies bearing any of several different DNA lesions, including one or more strand breaks and any one of many different possible large deletions. If the mtDNA is also incubated with Fpg to break the DNA at oxidative damage sites (8-oxo-G lesions) prior to beginning the PCR, then those lesions should also contribute to the total qPCR signal for mtDNA damage provided by this assay. (For assays of mtDNA-specific damage, care in primer design should be taken to ensure that P1 + P2 cannot amplify from nuclear mitochondrial-like pseudogenes, a.k.a. numts [18].)

In conclusion, this Primer Extension Blockade Enabled qPCR (PEBE-qPCR) method, a new addition to the already vast collection of assays in the PCR-toolkit, may facilitate many high-throughput, low-cost qPCR-based investigations in biology and medicine.

## Methods

### Human DNA samples

The human DNA sample used to develop the PEBE-qPCR 45S rDNA promoter assay is a pool of equal amounts of purified whole blood DNA from eight healthy Caucasian individuals aged 65 years or older from the general Utah population. DNAs were extracted using the Gentra Puregene Blood Kit (https://www.qiagen.com/us/shop/sample-technologies/dna/genomic-dna/gentra-puregene-blood-kit/#orderinginformation), dissolved in 10 mM Tris-Cl, 1 mM EDTA, pH 7.5 at 25°C, at a concentration of approximately 200 ng/μl, confirmed by agarose gel electrophoresis to consist of high molecular weight DNA with negligible degradation, and stored long-term at 4°C.

### Primers

The three oligonucleotide primers rpdd, rpd, and rpux (whose sequences are given in Fig. 2) were purchased from Integrated DNA Technologies, Inc. (idtdna.com), dissolved in 10 mM Tris-Cl, 0.1 mM EDTA, pH 8.0, to stock concentrations of 25 −100 μM, and stored at 4°C.

### Restriction endonucleases

MspI (at 100,000 units per ml) and HpaII (at 50,000 units per ml) were purchased from New England BioLabs, Inc. (neb.com) and stored at −20°C.

### Reaction mix composition

All chemical reactions in the assay were carried out in a closed-tube system containing 50 mM potassium acetate, 20 mM Tris-acetate, 10 mM magnesium acetate, and 100 μg/ml bovine serum albumin, pH 7.9 at 25°C (1X CutSmart Buffer, New England BioLabs, neb.com); 1X SYBR Green I (Invitrogen); 0.2 mM each dNTP (Life Technologies); 1M betaine (Life Technologies); and 1X Titanium Taq DNA Polymerase (Takara Bio USA). Concentrated stocks of these components were combined, and water added, to produce a 3X Master Mix which was stored in 1200 μl aliquots (enough for 360 ten μl reactions) at −20°C for months with no detectable loss of activity. (Many PCR applications require or benefit from digestion of the DNA with one or more restriction endonucleases prior to thermal cycling. NEB’s 1X CutSmart Buffer, which allows optimal restriction digestion by hundreds of different restriction enzymes, is also compatible with PCR when a DNA polymerase such as Titanium Taq, which tolerates high magnesium ion concentrations, is used.) Some reactions received MspI, at a final concentration of 5.2 units per 10 μl reaction; other reactions received HpaII at a final concentration of 2.6 units per 10 μl reaction. Final concentrations for the oligonucleotide primers were: rpdd, 500 nM; rpd, 250 nM; and rpux, 250 nM.

### Reaction setup and Standard Curves

All reactions were performed in 10 μl volumes, in Bio-Rad Hard-Shell® 384-Well PCR Plates, #HSP3805, on a Bio-Rad CFX384 Real-time Detection System. The SYBR Green I fluorescence data was analyzed with Bio-Rad’s CFX Manager software, File Version 3.1.1517.0823, using the Single Threshold C_q_ Determination Mode, with Auto Calculated Baseline Thresholds. To set up the Standard Curve reactions, 2-fold serial dilutions of the stock pooled DNA sample were prepared in Dilution Buffer (10 mM Tris-Cl, 0.1 mM EDTA, pH 8.0, with 100 ng/μl bovine serum albumin). Each reaction well received 5 μ l of 2X master mix containing the 3X Master Mix described above, restriction enzyme, primers, and water; and 5 μl of DNA or DNA-free Dilution Buffer. The input amounts of genomic DNA per reaction well for Standard Curve analyses were 1 ng, 2 ng, 4 ng, 8 ng, and 16 ng. Each reaction condition (+/-restriction enzyme, +/-various primers, with various DNA input amounts) was run in triplicate on each 384-well PCR plate. Microsoft Excel was used to generate the linear regression plots and r^2^ values shown in Fig. 3.

### Thermal profile and cycling

The thermal profile and cycling used is shown in Fig. 3a.

## Acknowledgements

We thank Richard Kerber, Elizabeth O’Brien, Chris Pappas, Jeff Stevens, and Mike Howard for helpful discussions.

## Availability of data

Figure 3, panels b, c, and d have associated raw data collected during the qPCR run on the Bio-Rad CFX384 Real-time Detection System, under the control of the Bio-Rad CFX Manager Software, and stored in a computer file. Copies of this file are available, without restriction, from Dr. Cawthon, upon request.

